# Early vegetative development and ontogenetic phase transitions in *Juglans neotropica* Diels: a BBCH-scale approach

**DOI:** 10.64898/2026.08.09.743696

**Authors:** Iván Delgado, María Alejandra Jaramillo, Fermín Rada, Pedro Jiménez

## Abstract

*Juglans neotropica* is an endangered South American walnut species for which standardized descriptions of early development are lacking, limiting its effective use in conservation and restoration programs. We developed a BBCH-scale description of early vegetative growth of *J. neotropica* based on observations under nursery and field conditions in Colombia. Three principal vegetative stages were described: seed germination (stage 0), leaf development (stage 1), and stem elongation (stage 3). Germination was hypogeal and occurred 30-140 d after sowing, occasionally extending to 180 d. During early growth, leaflet morphology, number, and architecture showed consistent and discrete changes between stages 1 and 3, including shifts in apex, base, margin type, and laminar shape. These modifications indicate that early development is organized into distinct ontogenetic phases rather than continuous variation, marking the transition from juvenile to vegetative adult stages. By providing a standardized, development-based framework independent of chronological age, this BBCH scale facilitates accurate identification and monitoring of seedlings in nursery production, restoration projects, and urban forestry programs. More broadly, this approach contributes to the characterization of ontogenetic phase transitions in tropical tree species and supports the use of development-based criteria for managing early establishment and performance.

## Introduction

*Juglans neotropica* Diels (Nogal) is the only walnut species native to the northern Andes, distributed across Colombia, Ecuador, Peru, and Venezuela (Manning 1960; Song et al. 2020). In Colombia, intensive logging has reduced 52% of its natural populations (Salinas and Cárdenas 2007), confining them mainly to protected areas such as Chingaza, Ukumari, and Cueva de los Guácharos National Parks (Ramírez and Kallarackal 2021; Ramírez 2022). Consequently, *J. neotropica* is currently listed as endangered in Colombia and protected under a national logging ban (Resolución No. 0316, 1974). Restoration efforts based on seed propagation and vegetative cuttings have been implemented (Botina 2011), and *J. neotropica* has been deployed as an urban tree in Bogotá (Ramírez and Kallarackal 2021; Ramírez, 2022). However, reliable methodological tools are needed to identify and monitor plants across developmental stages to support restoration efforts, particularly because chronological age alone is often a poor predictor of functional or physiological status in woody species.

Methodological tools for *J. neotropica*, such as standardized scales and botanical descriptions, are primarily focused on adult trees. Within the botanical descriptions, it has been established that adult trees can reach heights of up to 48 m and have gray bark marked with longitudinal grooves. Leaves are deciduous, alternate, and imparipinnate with opposite or sub-opposite leaflets. A characteristic aromatic compound of Juglandaceae, juglone, is produced by both the aerial and underground organs. This is a monoecious species bearing staminate catkins and pistillate flowers that produce black walnuts (Manning 1960; Toro-Vanegas and Roldán-Rojas 2018). These descriptions allow identification of adult trees but do not guide in determining vegetative developmental stages or recognizing juvenile plants. For understanding plant development and establishing management practices, standardized phenological scales are essential tools (Ramírez and Davenport 2020), as they enable consistent identification of functionally meaningful developmental stages across sites, nurseries, and monitoring programs. Ramírez and Kallarackal (2021) proposed a standardized description of *J. neotropica* vegetative and reproductive phenology; nevertheless, no framework exists for early developmental stages.

The *Biologische Bundesanstalt, Bundessortenamt, und Chemische Industrie* (BBCH) scale provides a numerical system for describing plant growth stages and has been widely applied to commercially important crops (Meier 2001). Its utility, however, extends beyond agriculture to conservation biology and ecology, particularly for woody tropical species in which chronological age is often a poor proxy for functional or developmental status, as morphology, physiology and performance vary consistently among ontogenetic stages (Poorter 2007; Wright et al. 2011). These ontogenetic transitions are common in neotropical canopy trees and have important consequences for establishment, survival and management, even when the precise duration of each phase varies among species (Soliz-Gamboa et al. 2011). The BBCH scale provides a standardized numerical system for describing plant development across the life cycle, from germination through reproductive stages (Meier 2001). Its modular structure allows descriptions to be tailored to specific developmental windows, enabling detailed characterization of early vegetative growth independently of later reproductive phases. By decoupling developmental state from plant age, the BBCH scale is particularly useful for woody species with prolonged juvenile phases. In *J. neotropica*, existing phenological frameworks and botanical descriptions focus almost exclusively on adult trees and reproductive stages, leaving early vegetative development poorly characterized despite its relevance for propagation and restoration programs (Ramírez and Kallarackal 2021). By decoupling developmental stage from chronological age, the BBCH framework offers a standardized approach for identifying functionally meaningful stages during early growth. Beyond species-specific relevance, early ontogenetic development in woody plants is increasingly recognized as structured by discrete phase shifts with functional consequences for plant performance (Poorter 2007; Wright et al. 2010), resource use and establishment success. However, standardized frameworks capturing these transitions remain scarce for tropical tree species, limiting comparisons across studies and their application in restoration contexts. In this study, we develop a BBCH scale for *J. neotropica* that describes early vegetative development from germination through stem elongation, providing a practical and transferable framework for characterizing ontogenetic phase transitions in the early development of tropical trees.

## Materials and methods

### Plant material and growth conditions

We collected 900 fruits from different mother trees in five municipalities along the Bogotá Savannah, Colombia: Bogotá, Cajicá, Chia, Subachoque, and Tabio. Fruits were pooled and mechanically scarified, and healthy, intact seeds were selected for further experiments. A total of 800 seeds were sown in 4-L containers filled with a 2:1 mixture of garden soil and rice husk. Containers were maintained in a nursery at Universidad Militar Nueva Granada, Cajicá, Colombia (4°55’11” N, 74°01’82” W) for fourteen months.

### BBCH scale development

The BBCH scale employs a decimal-based coding system to describe plant development across the entire life cycle, from germination to senescence. Principal growth stages are described by digits ranging from 0 to 9, with stages 0-4 describing germination and vegetative growth and stages 5-9 describing reproductive development. Within each principal stage, two-digit codes are used to distinguish secondary stages, while three-digit codes allow the definition of mesostages when finer developmental resolution is required (Meier 2001).

In this study, we applied the BBCH framework to characterize early vegetative development in *J. neotropica*, focusing on stages 0, 1, and 3, which correspond to germination, leaf development, and stem elongation, respectively. Stages 2 and 4, defined for monocotyledonous species, were not applicable and were therefore omitted. All descriptions followed BBCH phenological scale guidelines (Hess et al. 1997; Meier 2001), and morphological terminology was based on Simpson (2019).

Seeds were monitored weekly to document germination stages. For growth descriptions, two groups of plants were established as follows: Group 1: 400 individuals were used to describe stages 1 and 3. Group 2: 100 adult individuals from the Bogotá River restoration zone and urban trees in Cajicá were monitored during a year to validate and contextualize the phenological patterns observed in early stages. In total, 500 plants were considered for phenological descriptions.

## Results

We identified three principal stages of early vegetative growth: germination (Stage 0), leaf development (Stage 1), and stem elongation (Stage 3). Detailed stage descriptions are provided in Table 1 and illustrated in Figure 1, with morphological comparisons of leaves and leaflets between stages 1 and 3 shown in Figure 2. The combination of leaflet number, morphology, and recurring defoliation-regrowth cycles allowed clear delimitation of these stages as discrete ontogenetic phases rather that continuous variation.

**Table 1.**
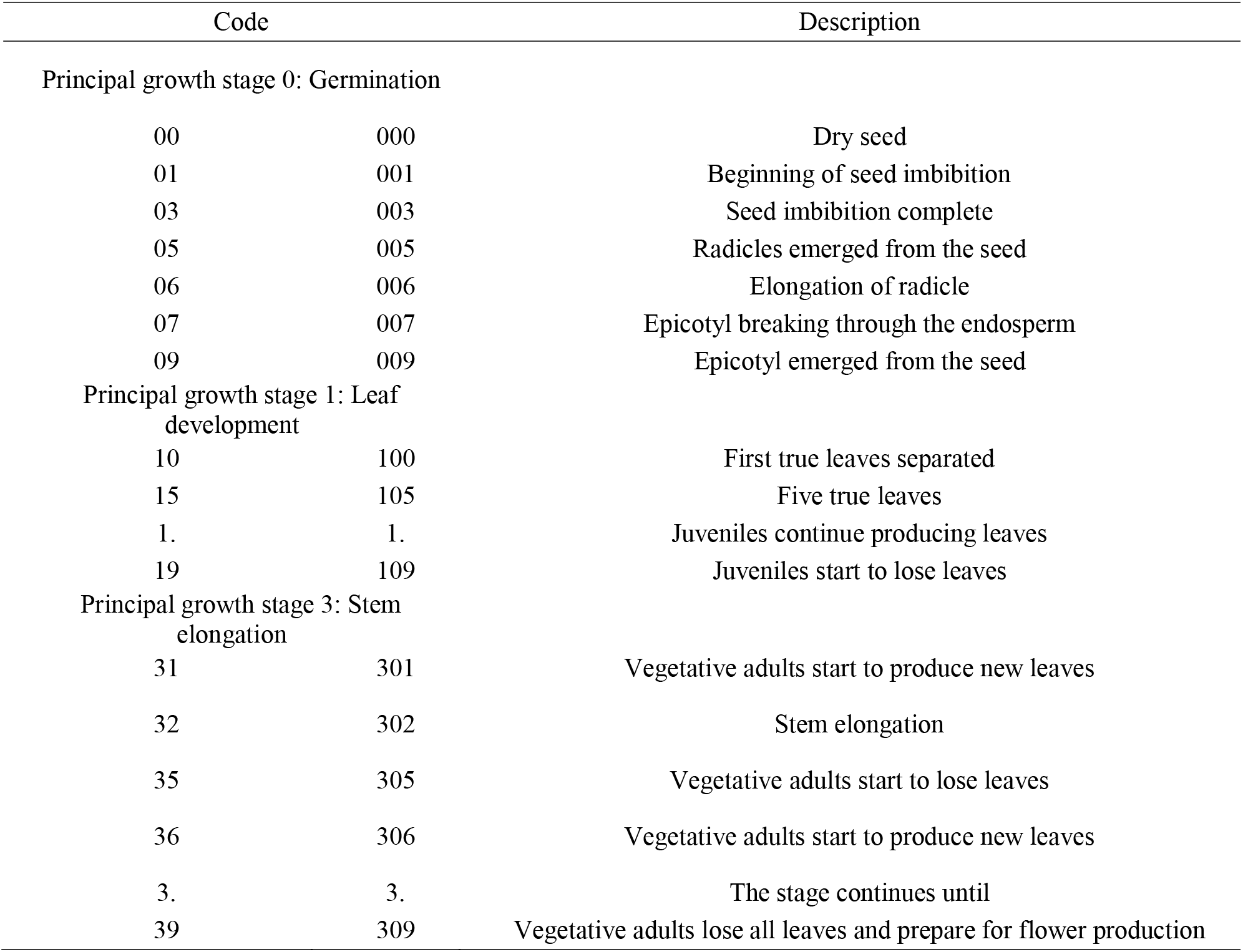
*Juglans neotropica* early vegetative growth according to the BBCH scale.

**Fig 1.**
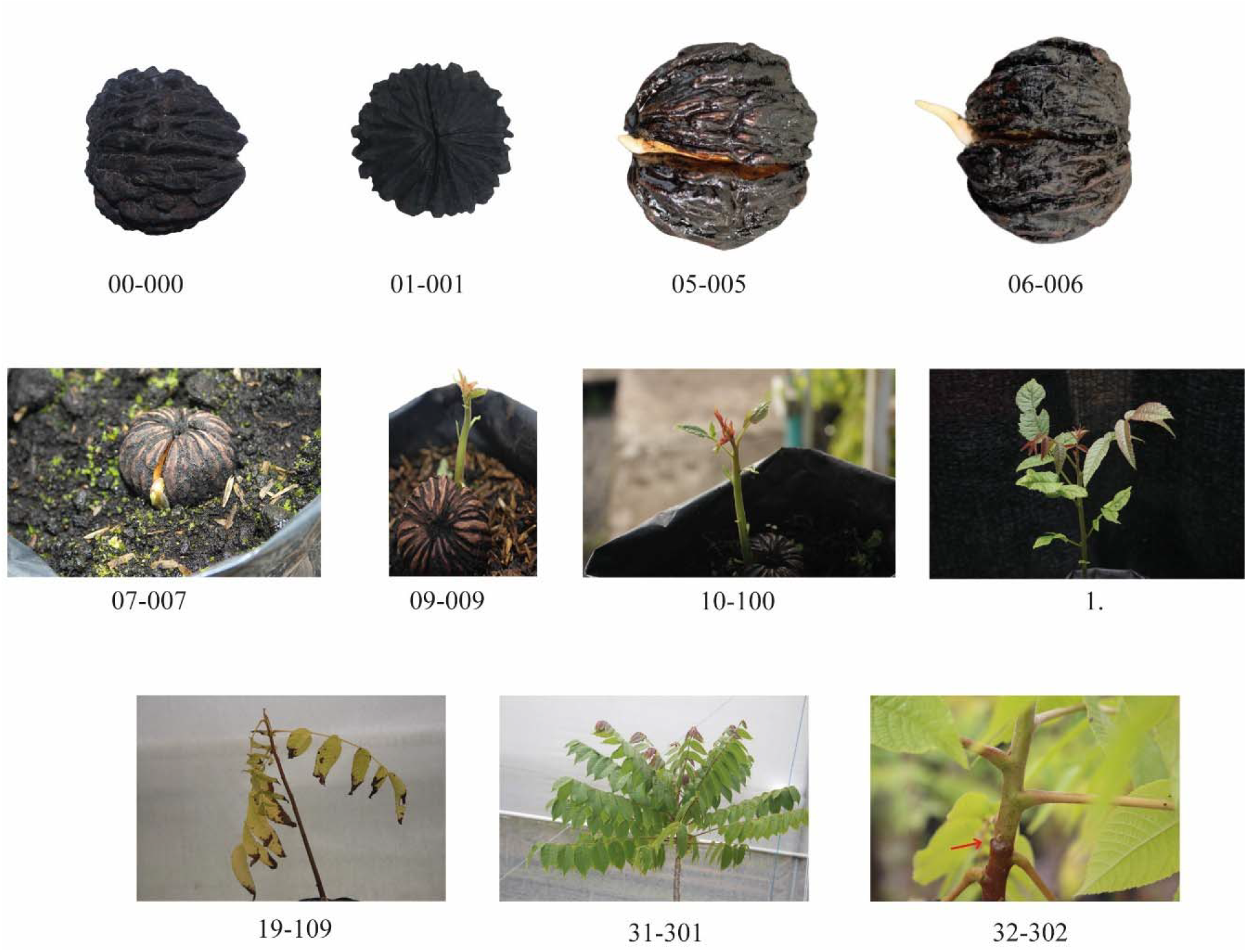
*Juglans neotropica* early vegetative growth stages. The red arrow indicates the boundary between the entirely hardened portion of the stem and the non-hardened portion

**Fig 2.**
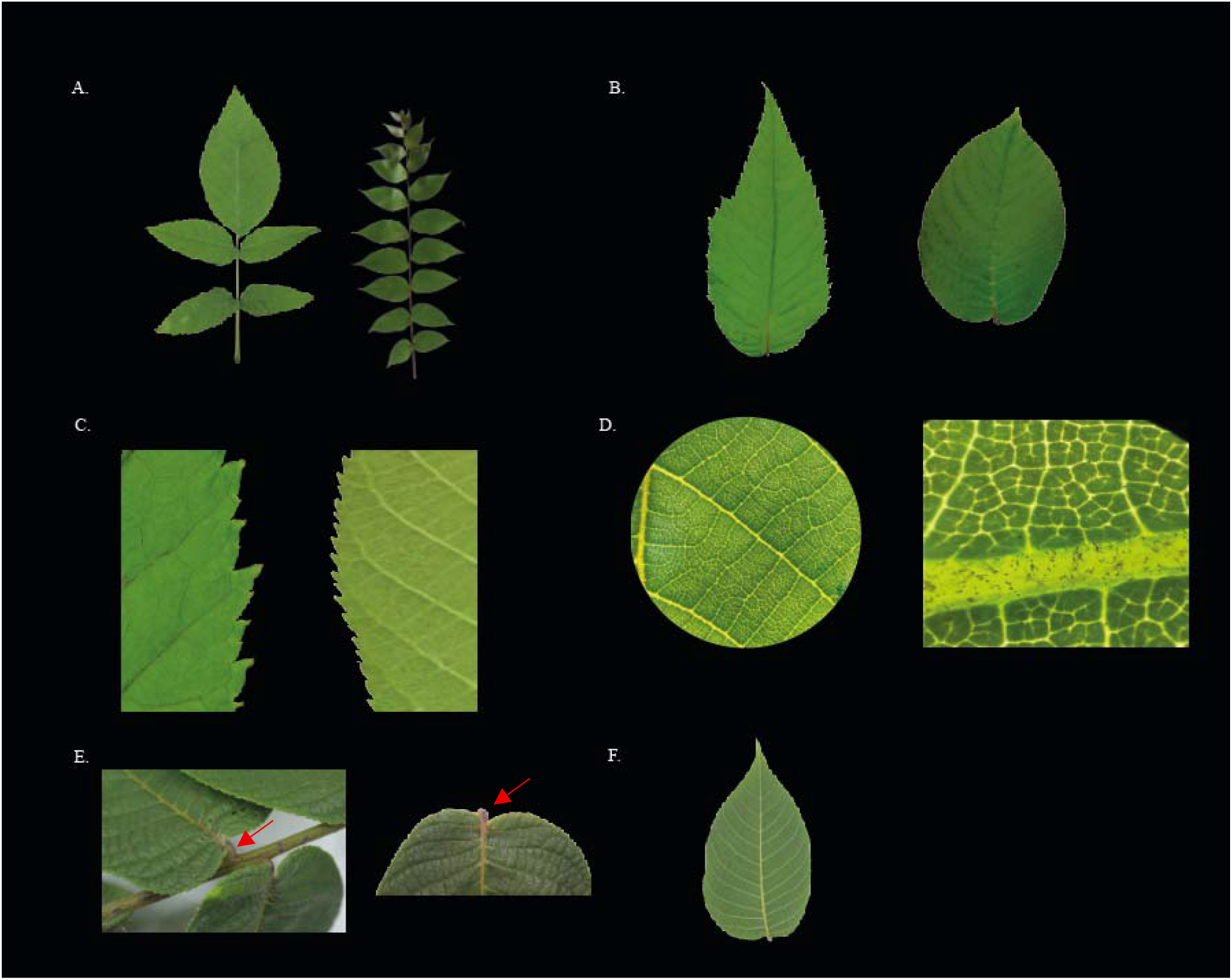
Differences in leaves and leaflets during stages 1 and 3 are shown from left to right. A Leaf: juvenile and vegetative adult. B Leaflet: juvenile and vegetative adult. C Leaflet edge: juvenile and vegetative adult. D Leaflet stereoscopic view: adaxial and abaxial surfaces, showing only stellate sessile trichomes along primary and secondary veins. E Petiolule: leaflet attached to the leaf and leaflet detached; the red arrow indicates the petiolule. F Leaflet venation

### Principal growth stage 0: Germination

Germination encompassed development from dry seed to epicotyl emergence (Table 1). The dry seed is a black walnut characterized by distinct distal and proximal poles and a hard shell bearing longitudinal ridges. At 5-7 d post-extraction, the valves at the distal pole opened, initiating imbibition. The valves closed after approximately 24 h, indicating complete imbibition (Figure 1).

The radicle protrusion occurred 20 d after sowing (physiological germination), followed by radicle elongation between days 20 and 30. Around four weeks after sowing, the epicotyl broke through the endosperm and emerged from the seed within 1-6 d after breaking through the endosperm (agronomic germination). Germination is hypogeal. Total germination time ranged from 30 to 140 d after sowing, although a second germination peak was observed as late as 180 d after sowing.

### Principal growth stage 1: Leaf development

This stage began with the emergence of the first true leaves and continued until seedlings experienced their first leaf loss (Table 1). Upon germination, plants typically produced 3-4 true leaves, each with a single leaflet resembling simple leaves. In some seedlings, this initial growth was absent, and plants developed 2-3 leaves with three leaflets each. During this stage, the stem transitioned from partially hardened to entirely hardened (Figure 1).

Leaves are compound, spirally alternate, and imparipinnate with 1-11 leaflets. Leaflets are sub-opposite near the petiole to opposite for those distally located, and present the following characteristics: elliptic shape, petiolulate (subsessile), cuneate base, acute apex, doubly serrate margins, pinnate simple and craspedodromous venation, membranous texture, abaxial surface is lighter green than adaxial side with glabrous interveinal spaces, and stellate sessile trichomes restricted to principal and secondary veins. At the end of this stage, foliage senescence occurs, and some individuals lose all foliage (Figure 2), marking the transition toward a new cycle of growth rather than plant mortality or stress-induced leaf loss.

### Principal growth stage 3: Stem elongation

This stage began when plants produced new leaves (Table 1). As the stem elongated, a distinct demarcation became visible between the entirely hardened basal portion and the non-hardened apical portion. Subsequently, plants underwent complete defoliation, followed by new leaf production and continued stem elongation. This recurring pattern of defoliation, renewed leaf production, and stem elongation constitutes a characteristic ontogenetic cycle and repeats several times before the plant reaches reproductive maturity. Each new leaf production cycle adds a pair of leaflets until the plant reaches the reproductive stage, and no more leaflets are added. Stem bifurcation can also initiate during this stage (Figure 1).

Leaves in stage 3 resembled those in stage 1 but contained 9 to 27 leaflets. While leaflet arrangement and venation patterns remained consistent, morphology differed: ovate (vs. elliptic) shape, truncate (vs. cuneate) base, acuminate (vs. acute) apex, serrulate (vs. doubly serrate) margins, and mesophytic (vs. membranous) texture. The abaxial side is lighter green than the adaxial side; it has glabrous interveinal spaces with stellate trichomes on the veins (Figure 2).

## Discussion

Several botanical descriptions of *J. neotropica* exist; however, most have focused exclusively on adult tree characteristics (Gómez et al. 2013; Manning 1960; Medina et al. 2021). A previous study by Ramirez and Kallarackal (2021) provided an extensive description of the phenological stages of *J. neotropica* adults using a BBCH scale. Here, we propose a BBCH scale for early growth in *J. neotropica*.

Postembryonic development in plants is divided into consecutive phases: juvenile, adult vegetative, and adult reproductive. Phase changes involve changes in morphological characteristics and are particularly common in woody plants (Taiz et al. 2017). Chronological age or simple size metrics often provide poor proxies for functional performance in tropical trees, as trait-performance relationships shift markedly across ontogenetic stages (Poorter 2007; Umaña et al. 2025). Our results documented morphological differences in leaflet apices, bases, edges, laminar shape, and number during the phase change from stage 1 to stage 3. Based on these observations, plants in stage 1 can be considered juveniles; whereas those in stage 3 should be regarded as adult vegetative (Taiz et al. 2017), since they resembled adult reproductive leaf characteristics (Manning 1960). These changes were not gradual but occurred as discrete shifts in leaflet morphology and architecture, suggesting the presence of ontogenetic thresholds rather than continuous developmental trajectories (Taiz et al. 2017). Such discontinuities are consistent with phase changes described in other woody species and may reflect coordinated changes in functional traits associated with resource acquisition and growth strategies (Poorter 2007).

Our observations on the number of leaflets during stage 3 (9-27 leaflets) differ from previous reports of 11-15 leaflets (Gómez et al. 2013), 15-19 leaflets (Manning 1960), and 16-23 leaflets (Medina et al. 2021). This variation may reflect differences in the phenological age of the sampled plants. As mentioned above, each new cycle of leaf production and stem elongation leads to the production of an additional pair of leaflets in each new leaf. The observed increase in leaflet number across successive growth cycles further supports the interpretation of early development as a structured ontogenetic process. Rather than reflecting plastic responses within a single developmental stage, these changes appear to represent progressive advancement along a developmental trajectory culminating in the vegetative adult phase (Taiz et al. 2017). This pattern may be common among long-lived tropical trees, where repeated cycles of leaf production and stem elongation contribute to functional maturation over extended periods (Poorter 2007). Additionally, Manning (1960) described leaflets as sessile; however, we consistently observed small but distinct petiolules across all developmental stages (Figure 2), indicating that leaflets should be classified as petiolulate (subsessile) rather than truly sessile.

This study provides the first standardized phenological description of early development in *J. neotropica*. The BBCH framework provides precise developmental terminology for research and monitoring programs of *J. neotropica*. This scale highlights morphological modifications associated with developmental phases and enables accurate identification of these phases.

Beyond its descriptive value, the BBCH scale proposed here has direct applicability in conservation and restoration programs. Standardized identification of early developmental stages allows practitioners to evaluate seedling performance, survival and establishment success based on ontogenetic status and not chronological age, which is particularly relevant for long-lived tropical trees (Poorter 2007). Phenological standardization also facilitates comparison across nurseries and restoration sites, synchronizes monitoring efforts, and supports trait-based approaches increasingly used in restoration ecology (Lavorel et al. 2007). By enabling unambiguous recognition of juvenile and vegetative adult phases, this framework supports informed selection of planting material, adaptive management, and long-term monitoring of *J. neotropica* populations in restoration and urban forestry contexts. Beyond *J. neotropica*, the framework proposed here highlights the importance of incorporating ontogenetic criteria into the study and management of tropical tree species. Standardized developmental scales such as BBCH offer a robust approach to compare early growth across species and environments, facilitating trait-based restoration strategies and improving the reproducibility of nursery and field studies (Lavorel et al. 2007).

Published guidelines for *J. neotropica* nursery management remain limited and lack standardization. Existing recommendations are largely based on arbitrary morphological thresholds rather than developmental criteria. For example, Ospina-Penagos et al. (2003) recommended sowing seeds in trays, transplanting seedlings to plastic bags when radicles emerge, and final field transplanting once plants reach approximately 25 cm in height, whereas Gómez et al. (2013) suggested field transplanting at a depth of 30-35 cm. We consider the tray-to-bag-to-field approach problematic, as repeated handling increases the risk of root damage and transplant stress, particularly in tree species with rapidly developing taproots (Rietveld 1989). Moreover, reliance on height-based thresholds ignores ontogenetic status and may compromise establishment success by decoupling management decisions from functional developmental stages.

Based on the BBCH framework proposed here, we recommend an alternative, development-based protocol: direct sowing into plastic bags with a capacity of ≥3 L to allow uninterrupted root development, followed by field transplanting at the end of stage 301. At this stage, plants possess photosynthetically active foliage and have completed initial stem lignification, conditions that may enhance post-transplant survival through improved root-soil contact and continued carbon assimilation during the establishment phase.

## Conclusions

This study provides the first standardized BBCH-scale description of early development in *Juglans neotropica* and highlights the role of discrete ontogenetic phase transitions in structuring early vegetative growth. By explicitly linking discrete morphological traits to ontogenetic stages, this framework enables consistent identification of juvenile and vegetative adult phases independent of chronological age and may be applicable to other tropical tree species with prolonged early developmental phases. The proposed BBCH scale offers a practical and transferable tool for nursery management, ecological research and restoration programs, and contributes to the development of standardized ontogenetic frameworks for tropical tree species.

## Statements and Declarations

### Funding

Resources for this research were used from the Phytopathology laboratory at the Universidad Militar Nueva Granada.

### Conflict of interest

The authors have no competing interests to declare that are relevant to the content of this article.

### Data availability

All relevant data and materials are included in the main manuscript.

## Contributions

ID and PJ were involved in methodological design and data analysis. ID conducted the research. PJ conceptualized and supervised the work. FR and MAJ were involved in the data analysis. ID wrote the first draft of the manuscript, and all authors were involved in review and editing. All authors accepted the final version of this manuscript.

## Notes

### Competing Interest Statement

The authors have declared no competing interest.

